# LPP3 mediates self-generation of chemotactic LPA gradients by melanoma cells

**DOI:** 10.1101/153510

**Authors:** Olivia Susanto, Yvette W.H. Koh, Nick Morrice, Sergey Tumanov, Peter A. Thomason, Matthew Nielson, Luke Tweedy, Andrew J. Muinonen-Martin, Jurre J. Kamphorst, Gillian M. Mackay, Robert H. Insall

## Abstract

Melanoma cells steer out of tumours using self-generated lysophosphatidic acid (LPA) gradients. The cells break down LPA, which is present at high levels around the tumours, creating a dynamic gradient that is low in the tumour and high outside. They then also migrate up this gradient, creating a complex and evolving outward chemotactic stimulus. Here we introduce a new assay for self-generated chemotaxis, and show that raising LPA levels causes a delay in migration rather than loss of chemotactic efficiency. Knockdown of the lipid phosphatase LPP3 - but not its homologues LPP1 or LPP2 - diminishes the cell’s ability to break down LPA. This is specific for chemotactically active LPAs, such as the 18:1 and 20:4 species. Inhibition of autotaxin-mediated LPA production does not diminish outward chemotaxis, but loss of LPP3-mediated LPA breakdown blocks it. Similarly, in both 2D and 3D invasion assays, knockdown of LPP3 diminishes melanoma cells’ ability to invade. Our results demonstrate that LPP3 is the key enzyme in melanoma cells’ breakdown of LPA, and confirm the importance of attractant breakdown in LPAmediated cell steering.

## Introduction

Malignant melanoma is the most aggressive form of skin cancer, accounting for the majority of skin cancer-related deaths [1]. Melanoma is highly treatable if diagnosed in its early stages while still localised, however patient prognosis worsens significantly once metastasis has occurred. Melanoma is often classified according to its localisation and growth characteristics, transitioning from early “radial growth phase” (RGP) plaques localised in the epidermis, to proliferative “vertical growth phase” (VGP) tumours which can break through the basement membrane into the dermis, increasing the likelihood of metastasis [2, 3]. The factors involved in bringing about melanoma metastasis are still not completely understood. Changes in cell-cell adhesion due to alterations in cadherin expression, and other changes in the gene expression programme related to epithelial-mesenchymal transition (EMT), may play a part by altering the cytoskeletal, invasive and migratory properties of cells and weakening cell-cell junctions [4]. However, in the case of melanoma, other drivers of metastasis are thought to be particularly important. One such driver is chemotaxis, in which cells migrate up gradients of extracellular signals. Yet the signals driving melanoma cell migration out of the primary tumour are poorly characterised. One signal that may contribute to melanoma metastasis is the phospholipid signalling molecule lysophosphatidic acid (LPA), which was recently discovered to be a strong chemoattractant for melanoma cells [5].

LPA has been implicated in a wide range of signalling pathways and cellular processes including proliferation, migration, survival and adhesion. It comprises a phosphate head group, glycerol backbone and acyl chain of varied length and saturation, characteristics which control biological activity [6]. Longer acyl chains and, in particular, unsaturated C:C bonds within the acyl group correlate with greater activity; short acyl chains and complete saturation make the LPA almost inactive [6]. The 18:1, 18:2 and 20:4 species (18 or 20 carbons in the acyl chain and 1, 2 or 4 unsaturated bonds) are thought to be the most active *in vivo*. LPA is found both intracellularly and extracellularly in tissues, and is also a major component of serum [7-9]. Extracellular LPA may be generated from lysophosphatidylcholine (LPC) *via* autotaxin, or converted from phosphatidic acid by membrane-bound phospholipase A1. However, source-sink gradients, in which one cell type produces LPA and another removes it to set up a chemotactic gradient, have not been found.

LPA mediates its effects through a family of seven-transmembrane G-protein coupled receptors. 6 LPA receptors have been identified (LPAR1-6), which bind LPA with varying affinity and interact with different combinations of the four Gα proteins Gα_12/13_, Gα_q/11_, Gα_i/o_ or Gα_s_ [9, 10]. LPA is broken down by a family of lipid phosphate phosphatases (LPPs). The LPPs are transmembrane enzymes located at the plasma membrane with an extracellular active site, and also on intracellular membranes. They dephosphorylate LPA into biologically inactive monoacylglycerol (MAG). There are three known mammalian LPPs (LPP1-3) and an additional splice variant of LPP1 [11]. LPP1 and PPAP2A are synonymous, LPP3 is also known as PPAP2B, while PPAP2C describes LPP2. This paper focuses on LPP3/PPAP2B, which will be simply described as “LPP3”.

LPA degradation by melanoma cells is of particular interest, as they appear to be able to form gradients with a low LPA concentration within the tumour and high LPA outside [5], thus setting up a “self-generated” gradient directing cells outwards from the primary tumour [12]. While LPA is a well-known inducer of cell motility and the role of LPA signalling in ovarian cancer is well studied [13, 14], it has only recently been shown to be a uniquely powerful chemoattractant for melanoma cells [5]. Thus LPA gradients could facilitate the exit and metastasis of melanoma cells from the tumour. Here we have investigated the mechanism by which melanoma cells break down LPA, demonstrating that LPP3 is crucial for the formation of self-generated LPA gradients and subsequent chemotaxis and invasion.

## Results

### Melanoma cells respond least efficiently to high concentrations of LPA

LPA is found in serum at varying concentrations of 0.1-10μM [7-9, 13, 15, 16], with an average of approximately 1μM, though this has been shown to increase under pathological conditions including various cancers. In contrast, the K_d_ of LPAR1, the most important LPA receptor, is approximately 10nM [17]. The concentration of LPA in serum – or even the 10% serum used in culture medium – is therefore saturating to most sensitive cells. As a significant difference in receptor occupancy across cells is required for directed chemotaxis, we investigated how melanoma cells are able to sense and respond to these high concentrations of LPA.

To test the response of melanoma cells to increasing concentrations of LPA, we used Insall chambers [5, 18] to visualise chemotaxis, in particular measuring evolution of chemotactic efficiency over time. The use of direct-viewing chambers allowed us to quantify chemotaxis in detail while cells migrated. One straightforward output is the chemotactic index, described by the cosine of the angle between the gradient and the cells’ migration (cos θ); positive chemotactic indices indicate migration up-gradient, with a value of >0.1 indicating significant chemotaxis.

When imposing an external chemotactic gradient across the cells, we found that increasing concentrations of LPA did not increase the efficiency of directional migration of WM239A metastatic melanoma cells (Figure 1). Although the WM239A cells were able to migrate towards LPA at all concentrations between 100nM-10μM, (Figure 1A), the efficiency of chemotaxis evolved differently as the concentration rose. With an initial concentration of 100nM LPA, cells were immediately chemotactic (Figure 1Bi), with a mean chemotactic index of 0.22 (±0.06 SEM) (Figure 1Bii). This dropped over time, likely reflecting the degradation of LPA. When using 1μM LPA, the initial level of chemotaxis was slightly lower (0.18 ±0.2) (Figure 1Bi), but increased to 0.23 (±0.3) by 18-24 hours. In contrast, at high concentrations of LPA (10μM), chemotaxis was initially negligible, increasing over 24 hours. Cells never reached the same chemotactic index as at lower concentrations of LPA. These results support the hypothesis that cells’ receptors are saturated at high concentrations of LPA, making it impossible for cells to perceive local changes, but they eventually break down LPA to sub-saturating levels that allow them to sense a gradient.

**Figure 1.**
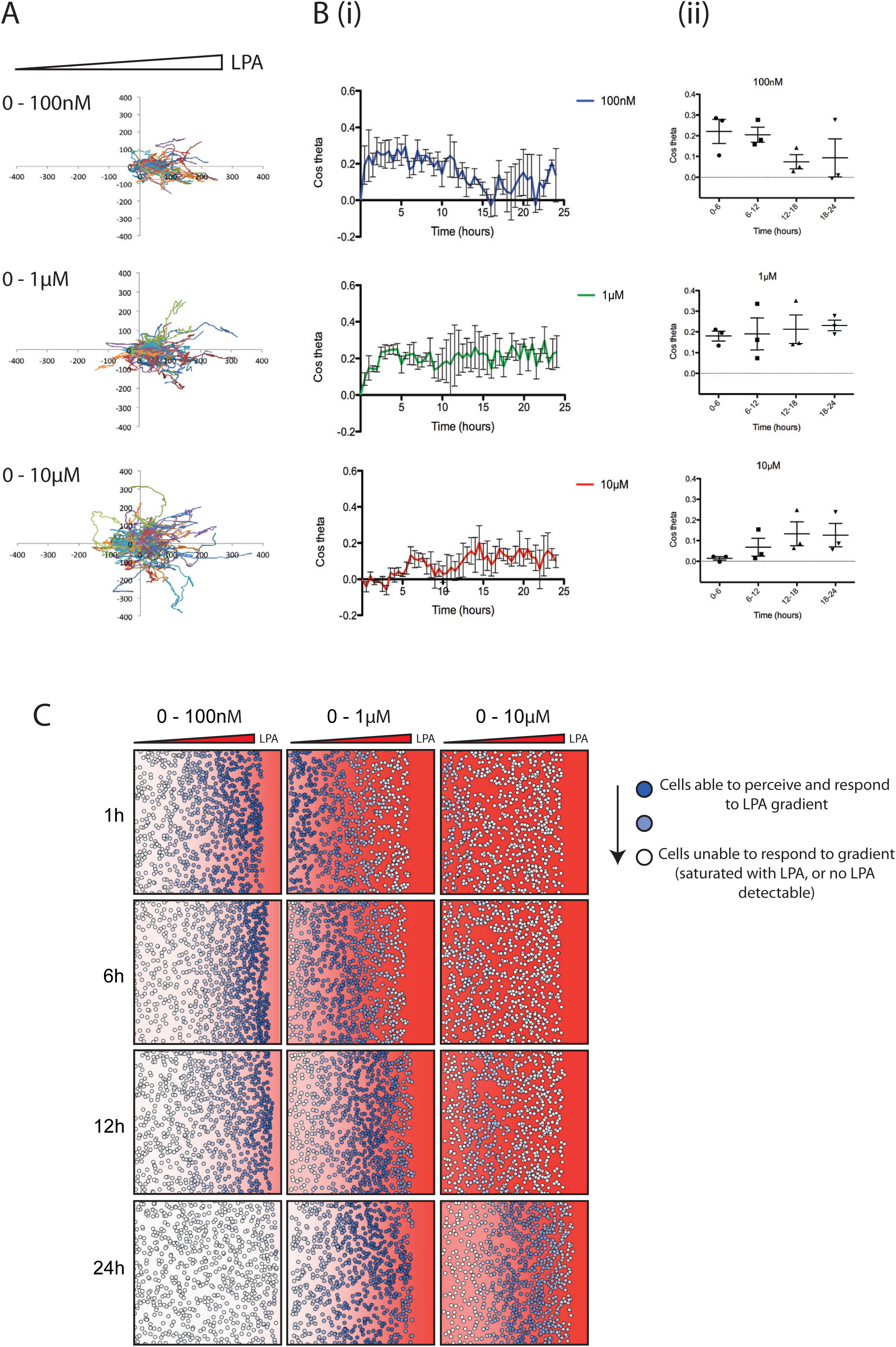
Higher LPA concentrations delay melanoma cell chemotaxis. (A) Representative spider plots showing cell tracks of WM239A metastatic melanoma cells moving from serum-free media (on the left) towards 100nM, 1μM or 10μM 18:1 LPA (on the right) in an Insall chemotaxis chamber. Axis scale is in μm. (B) Chemotactic response of WM239A cells over time, in the presence of increasing LPA concentration gradients. The mean (±SEM) chemotactic index of cells is shown (i) over 24 hours and (ii) across 6-hour time increments (n>100 cells for each condition, 3 independent experiments). (C) Representative snapshots over time from a mathematical model simulating cells locally degrading and responding to different concentrations of a freely diffusing chemoattractant (with similar kinetics and receptor affinities to LPA/LPAR). The red background represents the LPA concentration within the gradient and corresponding level of LPAR saturation. Solid red indicates a concentration of 1μM LPA or higher, and thus high receptor saturation. The colour fades linearly to white, which indicates a concentration of 0μM LPA, and thus no receptor saturation. Circles in dark blue represent cells able to perceive the LPA gradient, while circles fading to white represent cells increasingly unable to perceive the LPA gradient, either because no LPA is present or because too much LPA is present, oversaturating their receptors.

It is not currently technically possible to measure dynamic spatial LPA gradients directly. We therefore used a mathematical model that accurately represents the way cells modify gradients while responding to them [19] to further assess melanoma cell responses to varying levels of LPA. Figure 1C shows such a simulation of cells in different LPA gradients. At low initial LPA concentrations (0-100nM), most cells at the front respond at first as they are experiencing the steepest resolvable LPA gradient, however behind this front there is very little LPA present for cells to respond to. Over time the cells at the front also stop chemotaxing as the LPA is locally degraded. At intermediate concentrations (0-1μM), cells at the front are totally saturated and few initially respond. However, chemotaxis increases as the LPA is degraded to a level cells can respond to. At the highest concentrations (0-10μM), chemotaxis only starts once the majority of LPA is broken down, which takes at least 12hrs; even after this time, receptor saturation limits cells’ ability to perceive a gradient. The close agreement between model and measured data supports our hypothesis that changes in chemotaxis over time are mediated by the melanoma cells breaking down LPA as they respond to it.

### LPP3 is required for LPA breakdown by melanoma cells

Having demonstrated that the saturating LPA found in serum must be broken down in order to stimulate melanoma cell chemotaxis, we determined which enzymes from melanoma cells degrade LPA. We used siRNA to knock down expression of genes encoding each relevant mammalian LPP - LPP1, LPP2 and LPP3. Robust and specific knockdown was obtained in WM239A melanoma cells (Figure 2Ai). Next, LPA-containing medium was incubated for 24 hours with WM2349A cells, which were untreated or transfected with either a scrambled non-targeting siRNA or siRNAs targeting one LPP gene. The medium was analysed by mass spectrometry for LPA content, with the synthetic 17:0 form added during processing as an internal control. There was no significant difference between the different conditions in degradation of the less biologically active 18:0 form of LPA (Figure 2Aii). However, LPP3 knockdown resulted in significantly higher levels of 18:1 and 20:4 LPAs (Figure 2Aii), which are the biologically active forms responsible for signalling [6]. Over 60 hours, we found a clear delay in the rate of 18:1 and 20:4 LPA degradation when LPP3 was knocked down (Figure 2B).

**Figure 2.**
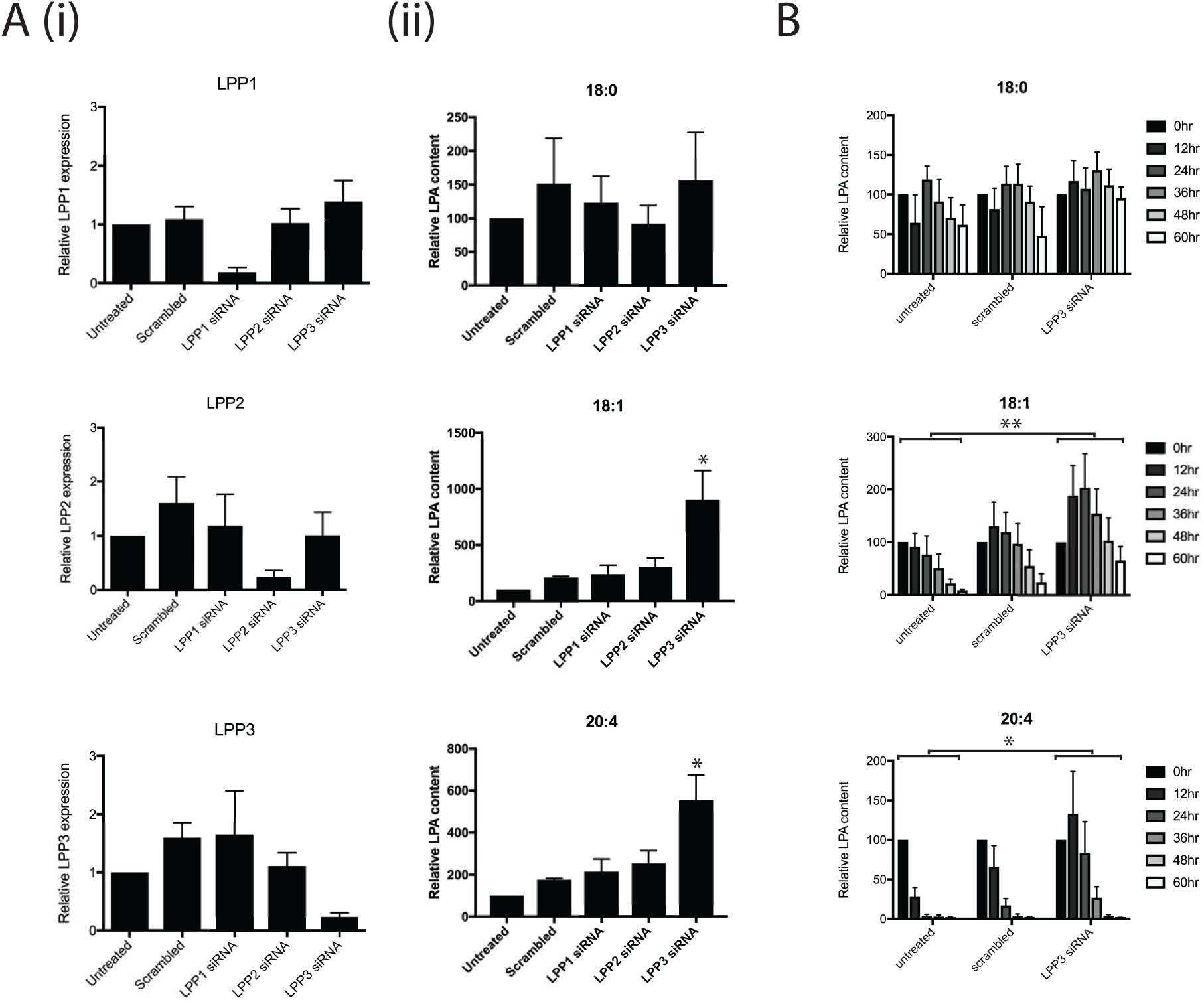
LPP3 is required for LPA breakdown by melanoma cells. (A) siRNA knockdown of the LPA breakdown enzymes LPP1, LPP2, LPP3 or a scrambled control, in WM239A melanoma cells. (i) Representative qPCR confirming knockdown of gene expression. (ii) Mass spectrometry quantifying relative LPA content in media cultured for 24 hours with WM239A cells. Different forms of LPA are shown: (top to bottom) 18:0, 18:1 and 20:4. (n=4, *p0.05 for LPP3 vs all other conditions, One-way ANOVA). (B) Timecourse of LPA breakdown by WM239A cells wither untreated or treated with control siRNA or siRNA targeting LPP3 alone. (n=4, *p_≤_0.05; *p<0.01 for untreated vs PPAP2B, Two-way ANOVA comparing the area under the curve)

A slight initial increase in 18:1 and 20:4 LPA at earlier timepoints suggests there is a background level of LPA production that is eventually normally compensated for by LPP3-mediated LPA breakdown. The addition of the autotaxin inhibitor HA130 leads to decreased LPA levels after 24 hours (Supplementary Figure 1), confirming that autotaxin can mediate further production of LPA over the saturating initial level.

Together, these results demonstrate that LPP3 is the key enzyme for degradation of the biologically active forms of LPA in melanoma cells.

### An assay to determine the role of LPA breakdown in self-generated chemotaxis

To investigate self-generated chemotaxis of melanoma cells towards LPA, we adapted the Insall chamber chemotaxis assay [20] (Figure 3Ai). Rather than imposing a gradient by adding serum-free and serum-containing medium to opposite wells, we added serum-containing medium to both to create an initially uniform environment. Since the viewing bridge is only 20μm high, the local volume of liquid is restricted, and the absolute amount of LPA is very small. However, the wells are far deeper, and thus contain a larger reservoir of LPA, which is consequently depleted much more slowly. Thus, local degradation of LPA by cells on the bridge would be expected to create a double outward-facing gradient, where cells would be attracted to LPA diffusing from the nearest well (Figure 3Aii). As predicted, in this assay cells initially positioned on the left side of the bridge (designated the “left cluster”) generally moved further to the left (Figure 3Bi-iii) and cells on the right (“right cluster”) moved rightwards (Figure 3Biv-vi, Supplementary movie 1). This effect was significant for both clusters (Rayleigh test for directionality; *p*<10^-9^), (Figure 3Bi, iv). Furthermore, directionality of cells in both clusters increased over time, from a slightly positive (rightwards) initial value to a substantial chemotactic index after 6 hours for the right cluster (Figure 3Bvi), and corresponding negative (leftwards) values for the left cluster (Figure 3Biii). We hypothesised that this delay was due to the time taken to break down LPA and create a gradient.

**Figure 3.**
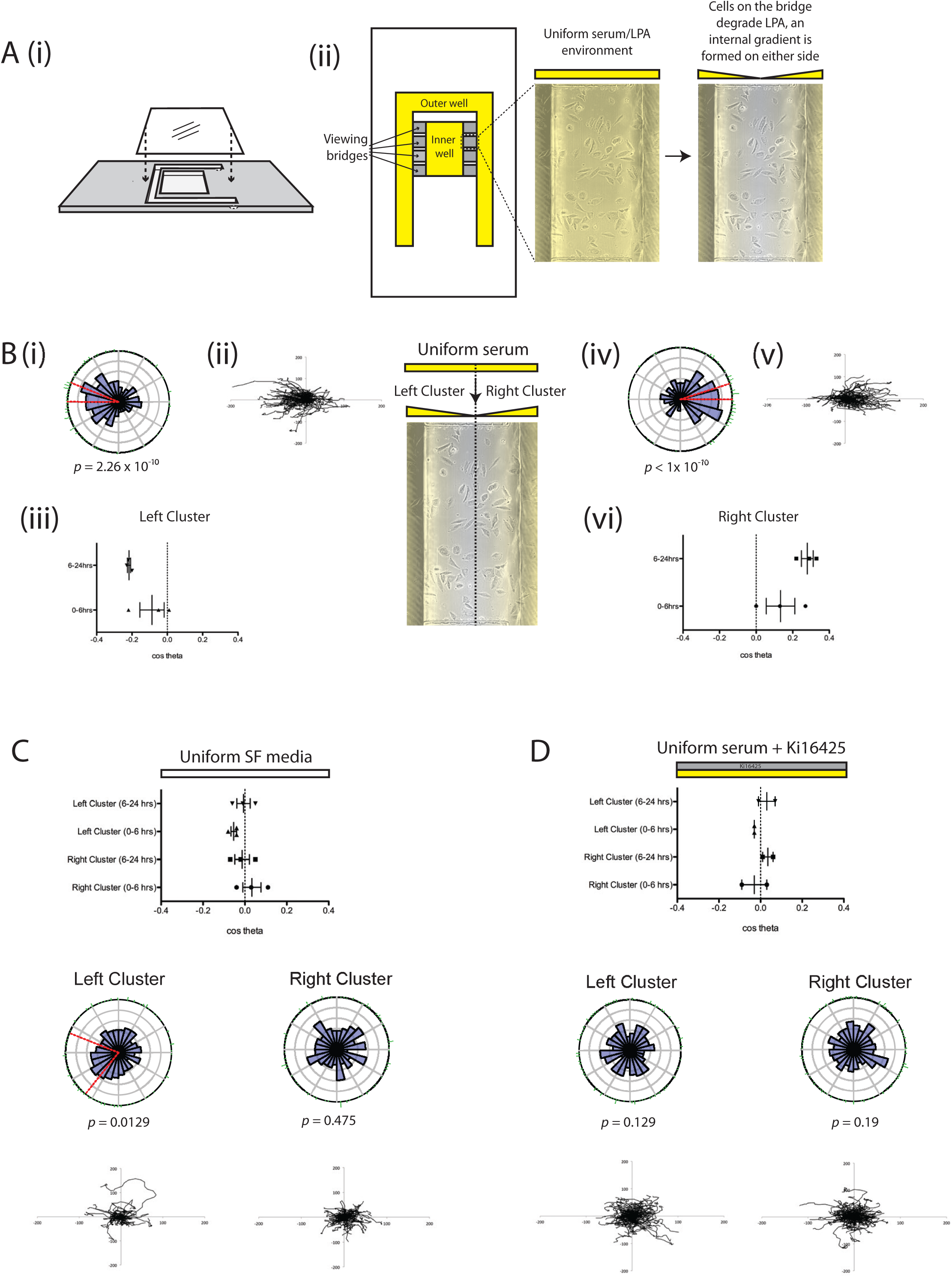
Spread assay to assess LPA breakdown and chemotaxis. (A) (i) Side-view of an Insall chemotaxis chamber, showing inner and outer wells and positioning of the coverslip with cells adhered to it (adapted from [33]). (ii) Bird’s eye view of an Insall chamber, indicating the position of the viewing bridges (in yellow). (B) A uniform serum environment produces differential directional movement of the cells on the left (Left cluster; i-iii) and right (Right cluster; iv-vi) of the bridge, as quantified by mean (±SEM) chemotactic index (iii, vi) or shown in representative rose plots with Rayleigh tests for directionality (i, iv) and the corresponding spider plots (ii, v) (n>100 cells per experiment, 3 independent experiments). In conditions where there is a uniform serum-free environment (C) or uniform serum with the addition of 10μM Ki16425 (D) there is no significant difference in cell directionality between the left and right clusters (n>100 cells per experiment, 2-3 independent experiments).

To confirm that this effect was specific to LPA, not a random effect reflecting the proximity of cells to the edge of the bridge, we tested the movement of cells in uniform serum-free conditions (Figure 3C, Supplementary movie 2) and in a uniform serum-containing environment containing the LPA receptor antagonist Ki16425 (Figure 3D, Supplementary movie 3). In both cases, we observed no significant directional movement of cells in either cluster.

### LPP3 knockdown inhibits LPA breakdown-mediated chemotaxis

This assay was used to demonstrate that inhibiting LPA breakdown by LPP3 knockdown prevents the formation of outward-facing gradients, and thus prevents self-generated chemotaxis. We used siRNA to transiently knockdown LPP3 expression, utilising two individual oligos that consistently reduced LPP3 expression relative to scrambled controls (Supplementary Figure 2). Firstly we demonstrated that LPP3 knockdown itself did not affect the response of melanoma cells to LPA in an external serum gradient (Figure 4A). LPP3 siRNA-treated cells and scrambled-siRNA controls had similar mean chemotactic indices (0.195 and 0.15, respectively; Figure 4Ai), and both populations showed highly directed migration towards serum with Rayleigh *p*-values of <10^-10^ (Figure 4Aii). However, in the self-generated chemotaxis assay, LPP3 knockdown cells displayed obviously different behaviour from controls. Control cells produced gradients that triggered chemotaxis of left and right clusters in opposite directions (Figure 4Bi). In contrast, LPP3 knockdown caused a significant reduction in chemotaxis of both clusters, with virtually no directional migration (p<0.05; Figure 4Bii, Supplementary movie 4). This was confirmed using the second independent siRNA targeting LPP3 (p<0.01; Figure 4Biii).

**Figure 4.**
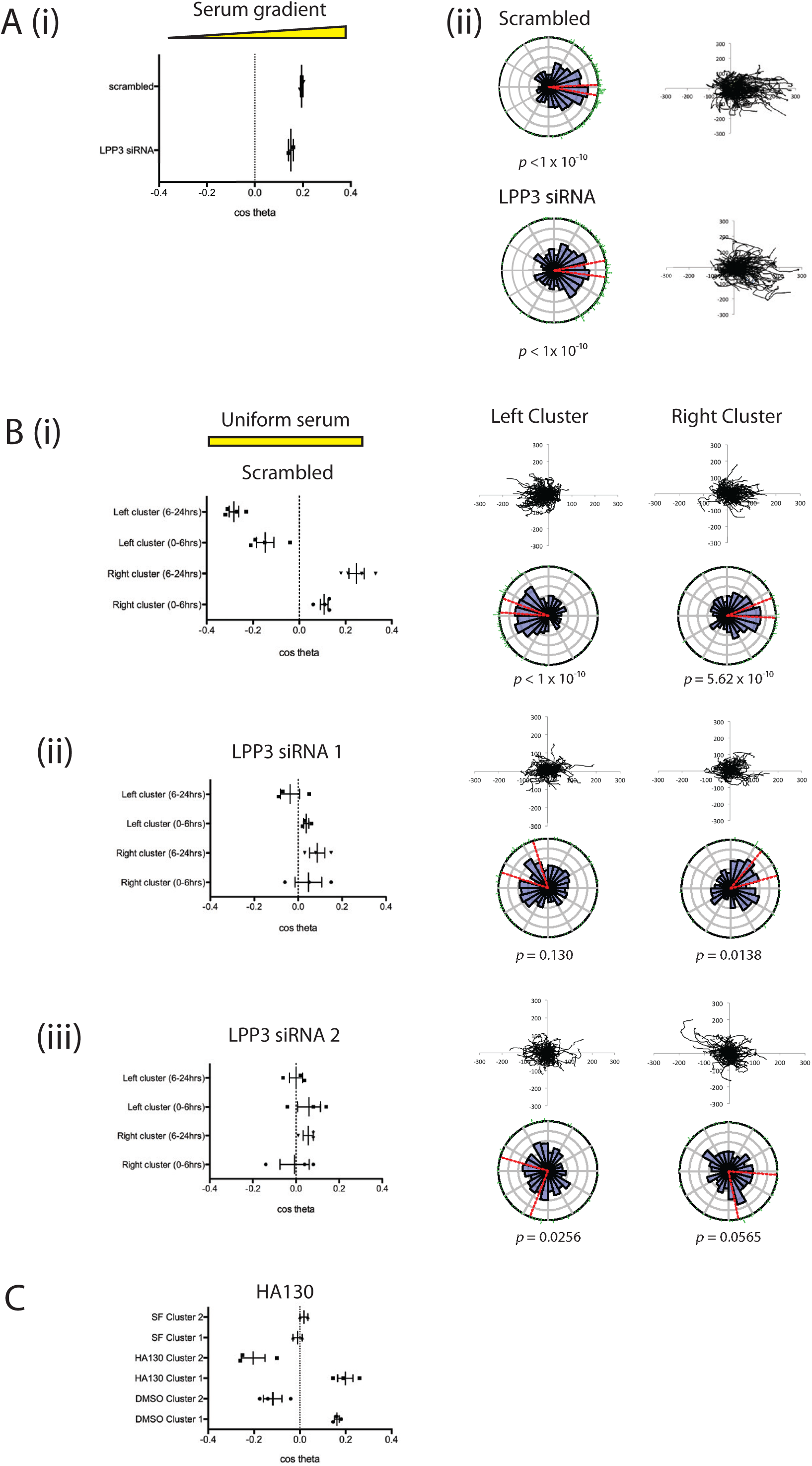
LPP3 knockdown inhibits chemotaxis in uniform serum. (A) LPP3 knockdown (LPP3 siRNA 1) does not inhibit chemotaxis of melanoma cells in the presence of a serum gradient, as assessed by (i) chemotactic index or (ii) rose plot and spider plot. (n>100 cells per experiment, 2 independent experiments). (B) LPP3 knockdown inhibits differential chemotaxis of melanoma cells in a uniform serum gradient, as shown when comparing (i) scrambled siRNA controls against cells targeted with two different siRNAs against LPP3 (ii, iii). (n>100 cells per experiment, 4 independent experiments. *p*<0.05 comparing clusters 1 and 2 of scrambled vs LPP3 siRNA 1 and 2, One-way ANOVA). (C) Chemotaxis of melanoma cells in a uniform serum gradient in the presence of either DMSO or 500nM HA130 to inhibit autotaxin function, or in a uniform serum free (SF) gradient. (n>100 cells per experiment, 3 independent experiments).

If LPA production via autotaxin were important in cell steering, we would expect the opposite results to what we observe. Wherever cell density was high, we would expect to see more LPA and therefore an attractive stimulus, so cells would tend to aggregate rather than dispersing outwards. However, since autotaxin is widely described as a promoter of metastasis, we directly tested its effects. We performed the assay for cell dispersal in the presence of the autotaxin inhibitor HA130 (Figure 4C, Supplementary movie 5). Inhibition of autotaxin did not inhibit differential chemotaxis of melanoma cells in this assay; if anything the inhibition of background LPA production enhanced chemotaxis in response to self-generated gradients. This demonstrates that degradation, rather than production of the chemoattractant LPA is the key process required for production of self-generated gradients and subsequent cell migration.

### LPP3, LPA and 3D invasion

We also investigated whether melanoma cell invasion involved self-generated gradients, and whether LPP3 was required to generate them. First we used circular invasion assays (CIAs). The CIA enables direct visualisation of cell migration in a manner similar to direct-view chambers, but in a setting that mimics invasion into 3D extracellular matrix, by forcing the cells to invade into Matrigel [21]. We used a different metastatic melanoma cell line, WM1158, as it displayed higher baseline levels of invasion. We observed that LPP3 knockdown resulted in reduced levels of invasion compared to controls (p<0.01; Figure 5A; Supplementary movie 6). As described earlier, this implies that outward-facing gradients of LPA are more important than the LPA concentration itself, and that LPP3 is instrumental in forming them.

**Figure 5.**
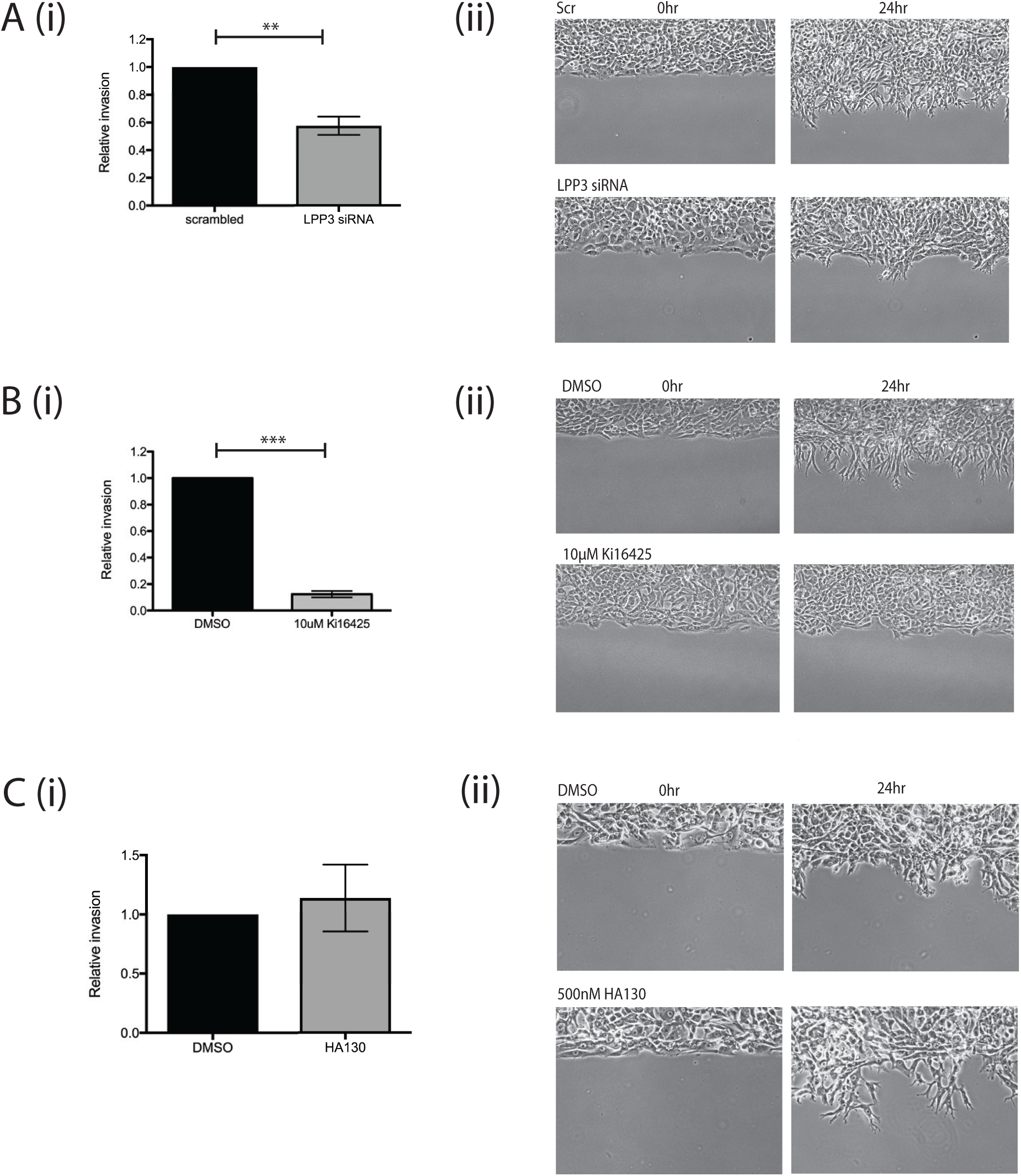
LPP3 knockdown inhibits melanoma cell invasion in 2D. (A) (i) Quantification of invasion of WM1158 metastatic melanoma cells transfected with either a scrambled control siRNA or siRNA targeting LPP3 (LPP3 siRNA 1) in a CIA. Mean +SEM shown (n=4 independent experiments, p<0.01, Student’s *t* test). (ii) Representative images of cell invasion at 0 and 24 hours. (B) (i) Quantification of invasion of WM1158 cells treated with DMSO control or 10μM Ki16425 in a CIA (n=3 independent experiments, p<0.0001, Student’s *t* test). (ii) Representative images of cell invasion at 0 and 24 hours. (C) (i) Circular invasion assay using WM1158 cells treated with DMSO control or 500nM HA130 (n=3 independent experiments). (ii) Representative images of cell invasion at 0 and 24 hours.

LPP3 has been reported to be important for cell-cell adhesion and ECM modulation by endothelial cells [22]. To verify that the reduction in melanoma cell invasion we measured was due to LPA gradient formation being blocked and not due to other LPP3-mediated interactions, we tested whether blocking LPA signalling specifically would inhibit invasion as well. Treatment of cells before and during the assay with Ki16425 produced a similar effect, markedly inhibiting invasion compared to controls (p<0.0001; Figure 5B; Supplementary movie 7). In contrast, addition of the autotaxin inhibitor had no effect on melanoma cell invasion into the matrix (Figure 5C; Supplementary movie 8). Thus, the formation of self-generated LPA gradients by LPP3 appears to be a key factor in the chemotaxis and invasion of melanoma cells.

Finally, we also examined spheroids to determine whether LPP3 expression and self-generated gradients were important for invasion in a 3D assay. We used the metastatic melanoma cell line WM852, which was the most clearly invasive line available in spheroid assays, treated with siRNAs targeting LPP3 or scrambled controls. Spheroids were imaged for 6 days following the addition of the invasion matrix, with the control spheroids producing many invasive protrusions outwards from the centre and displaying eventual dispersal of the cells from the central mass (Figure 6A). In comparison, the LPP3 knockdown spheroids displayed reduced invasion (Figure 6B). Although LPP3 siRNA knockdown was only transient and the effect dissipated by day 6 (Figure 6C), the effect was still potent enough to shorten and reduce the number of protrusions being generated, in addition to blocking cell dispersal outwards from the spheroid. Differences in proliferation appeared to have a minimal effect, as in the absence of invasion matrix the control spheroids were only slightly larger than the LPP3 knockdown spheroids (Figure 6D).

**Figure 6.**
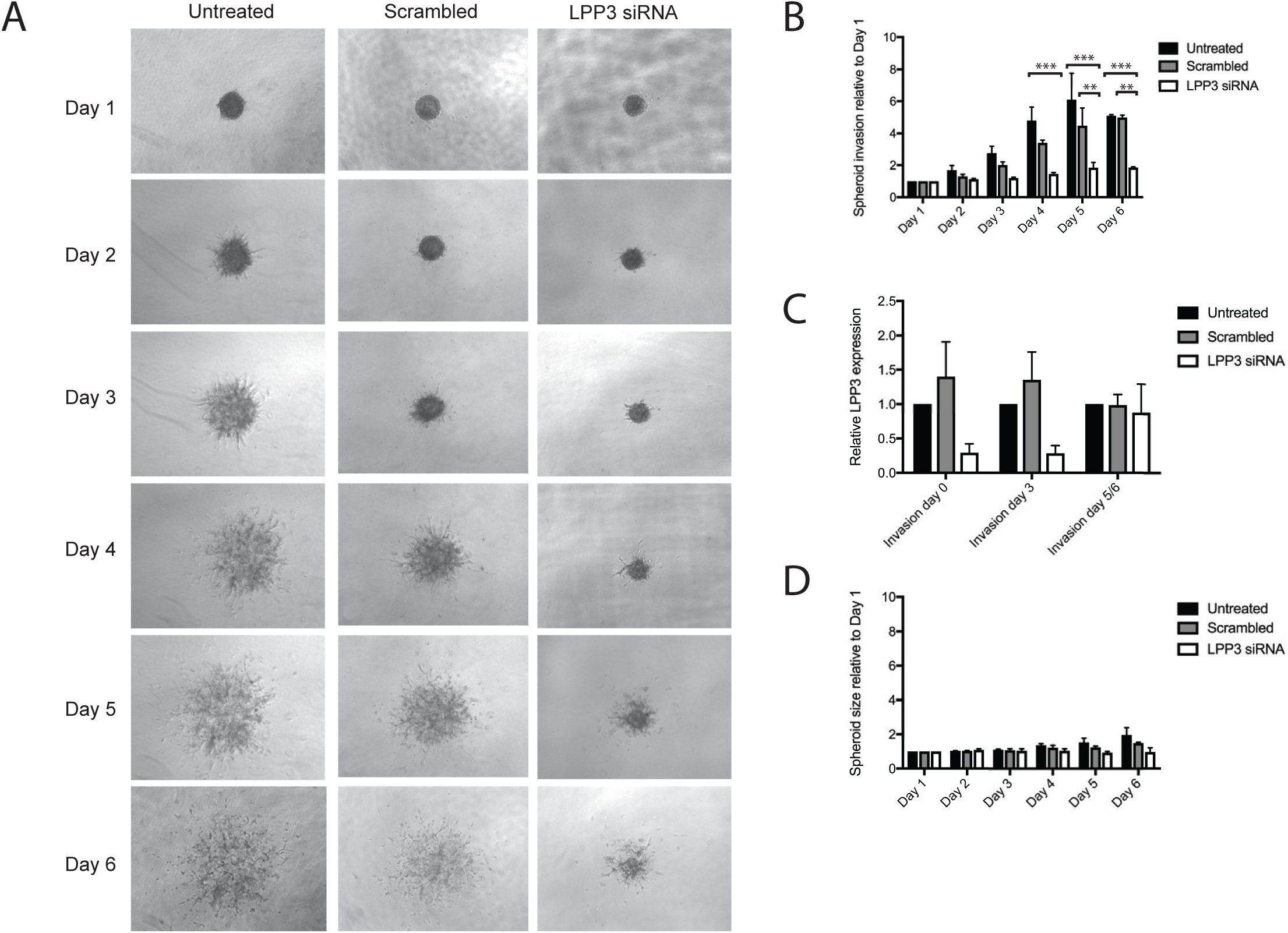
LPP3 knockdown inhibits melanoma cell invasion in 3D. 3D spheroid invasion assay using WM852 metastatic melanoma cells either left untreated or transfected with either a scrambled control siRNA or LPP3 siRNA 1. (A) Representative images of spheroid invasion over 6 days following the addition of invasion matrix. (B) Quantification of 3D invasion relative to spheroid size on Day 1. Mean +SEM shown (n=4 independent experiments, **p<0.01; ***p<0.001 Two-way ANOVA). (C) qPCR of *LPP3* expression in WM852 at Day 0, 3 and 5/6 following addition of invasion matrix (n=3 independent experiments). (D) Quantification of spheroid size relative to spheroids on Day 1, in the absence of any invasion matrix (n=3 independent experiments).

Overall, these results show that LPA is important in multiple assays for melanoma cell migration and invasion, and LPP3 plays a key part in these responses by shaping externally provided LPA into chemotactic gradients that steer individual cells away from any area with high density of cells.

## Discussion

We have previously demonstrated the efficiency of self-generated gradients as a driver of cell migration [19] and that self-generated LPA gradients are key factors in melanoma cell chemotaxis and thus metastasis [5]. Here, we have delineated the mechanism by which melanoma cells may be triggered to migrate away from the tumour *via* self-generated gradients.

LPA is ubiquitous in serum, and also found in healthy tissues. Serum LPA is often present at a high base concentration of 1μM (and significantly higher in some cancers). At this concentration, receptors at the front and rear of cells are all saturated, so cells are unable to detect LPA gradients and trigger chemotaxis. In order to read the gradient, cells need time to degrade the LPA to a level that does not saturate the LPA receptors, so that they can perceive spatial variations in LPA concentration as local differences in receptor occupancy. The higher the LPA concentration, the longer before the cells can respond. Similarly, the more cells available to degrade the LPA, the better they are able to respond [5]. Our data demonstrate that LPP3 is the key enzyme required for melanoma LPA breakdown. Furthermore, we have shown using various different metastatic melanoma cell lines in a range of 2D and 3D assays that LPP3 is essential for the production of a self-generated LPA gradient which can stimulate melanoma cell chemotaxis and invasion.

### LPP3 plays a non-redundant role in LPA breakdown

Although all three LPPs are thought to have similar catalytic activity, they appear to have non-redundant functions as evidenced by the different phenotypes observed in *Ppap2a* (LPP1), *Ppap2b* (LPP3) and *Ppap2c* (LPP2) knockout mice. While *Ppap2a* and *Ppap2c* knockout mice display only very mild phenotypes [23, 24], *Ppap2b* (LPP3) knockouts are embryonic lethal due to abnormal development of the vasculature [25]. This may indicate the need for self-generated LPA gradients in development; elsewhere, self-generated S1P gradients have been shown to be essential for T-cell function in the lymph system [27]. We also found that knockdown of LPP1 and LPP2 did not have any overt effect on LPA degradation, while LPP*3* knockdown resulted in initially increased levels of LPA and slower rates of breakdown over time. Thus, LPP3 appears to be the dominant lipid phosphatase required for LPA breakdown by melanoma cells.

### LPP3 is responsible for melanoma-mediated self-generated gradients

As there are no reagents currently available to directly visualise LPA degradation, we have developed a robust assay that can be used to examine and quantify cell responses to the formation of localised self-generated gradients. We have shown that after approximately 6 hours, melanoma cells in uniform serum-containing medium migrate towards the nearest LPA source. This effect is lost in serum-free medium, indicating that it is not merely an effect of random migration. Most importantly, LPP3 knockdown also abolished this effect, establishing the link between LPA breakdown and the ability of melanoma cells to form gradients in uniform serum. Although LPP3 is also capable of breaking down the bioactive lipid S1P, it is unlikely that S1P is involved in these self-generated gradients as the specific LPA receptor antagonist Ki16425 fully blocked chemotaxis.

There is a substantial literature implicating the phosphodiesterase autotaxin, which generates LPA from LPC, in cancer spread and chemotaxis [26, 27]. However, self-generated chemoattractant gradients generated by autotaxin would be expected to be high where cells were densest (at the centre of tumours), and therefore oppose cancer cell spreading and invasion. We have shown that in contrast to LPP3 knockdown, the addition of the autotaxin inhibitor HA130 does not abolish the divergent chemotaxis of melanoma cells in our assay, in fact the effect is slightly enhanced. This may be due to the elimination of a background level of LPA production, which could blur the effect of the self-generated gradient. Thus, this data confirms that it is the breakdown of LPA and formation of self-generated gradients that is the key factor responsible for driving melanoma cell migration, rather than autotaxin-mediated LPA production. As LPA has a split role – as well as being a chemoattractant, it is an exceptionally potent mitogen – it seems more likely that autotaxin’s principal role is to promote tumour growth. Breakdown of attractants, like the LPP3-mediated process we have described, generates gradients that lead away from the tumour and is a more likely driver of metastasis.

### A role for LPP3 in melanoma metastasis

While chemotaxis assays are useful for determining cellular responses to chemoattractants in 2D, they do not emulate the tissue environment faced by tumour cells when metastasising. Here we have used CIAs and 3D spheroid invasion assays to examine melanoma cells invading a matrix-filled environment. Although the serum-containing medium in a CIA/spheroid assay is not flow-restricted as in an Insall chamber, the LPA may be broken down locally within the matrigel component containing the cells, enabling self-generated LPA gradients to be maintained. Indeed, we found that LPP3 knockdown significantly inhibited LPA-dependent melanoma cell invasion, reinforcing the role of self-generated LPA gradients in facilitating melanoma metastasis.

These findings present an interesting potential target for anti-metastatic therapy when treating melanoma. Interestingly, LPP1 and LPP3 have been targeted in potential therapies for ovarian cancer [14, 28, 29]. However, these therapies aimed to increase LPP expression in order to reduce overall LPA levels, thereby diminishing its protumourigenic effects. In contrast, we would hypothesise that increasing LPP3 expression in melanoma cells would speed up their ability to create self-generated gradients, and thus accelerate migration out of the tumour. Thus, to diminish metastasis, LPP3 could potentially be neutralised to inhibit LPA breakdown around a tumour site.

In future, it will be interesting to test the universality of self-generated gradients in other processes and systems. LPP3 has previously been shown to be essential in depleting S1P within the thymus and spleen, setting up S1P gradients that stimulate lymphocyte migration, while its homolog has been shown to be involved in germ cell migration in *Drosophila* [30-32]. Thus, signalling lipid breakdown to form self-generated gradients may be common in both normal physiology and metastasis of various types of cancer, warranting further investigation into this simple yet powerful mechanism of cell steering.

## Methods

### Cell lines and reagents

WM239A, WM1158 and WM852 melanoma cells lines were obtained from the Wellcome Trust Functional Genomics Cell Bank (Biomedical Sciences Research Centre, St. George’s, University of London). Cells were cultured in RPMI medium (Invitrogen) supplemented with 2mM L-glutamine (Gibco) and 10% foetal bovine serum (FBS, PAA Labs).

1-oleoyl-2-hydroxy-*sn*-glycero-3-phosphate (18:1) and 1-arachidonoyl-2-hydroxy-*sn*glycero-3-phosphate (20:4) LPA was obtained from Avanti. FlexiTube GeneSolution sets of four siRNAs (Qiagen) were used to target LPP1, LPP2 and LPP3. LPP3 siRNA 1 was oligo 6 from the ON-TARGET plus siRNA range (catalogue no. J017312-06-0005; GE Dharmacon. siRNA sequence: GGGACUGUCUCGCGUAUCA), LPP3 siRNA 2 was oligo 7 from the FlexiTube siRNA range (catalogue no. SI03043761, Qiagen. siRNA sequence: AGCGATCGTCCCGGAGAGCAA), scrambled non-targeting siRNA control was AllStars negative control siRNA (Qiagen). Ki16425 (Cambridge Bio) was used at a final concentration of 10μM. HA130 (Echelon biosciences) was used at a final concentration of 500nM.

### Insall chamber chemotaxis assay

Cells were plated on fibronectin-coated glass coverslips at a density of 7x10^4^ cells/ml then serum starved overnight in serum-free RPMI medium prior to use as previously described [18, 33]. Serum-free (SF) medium comprised RPMI with 2mM L-glutamine, penicillin/streptomycin (Invitrogen) and 0.5% BSA (Sigma), while serum-containing medium contained an additional 10% FBS. LPA-containing medium comprised SF medium with LPA at the indicated final concentrations.

Insall chambers were imaged on a Nikon TE2000-E inverted timelapse microscope equipped with MetaMorph (Molecular Devices), a motorised stage (Prior Scientific, Cambridge UK) and Perfect Focus System enclosed in a plexiglass box humidified and heated to 37**°**C with 5% CO_2_. Each viewing bridge on the Insall chamber was imaged with a 10x objective every 30 minutes over 24 hours.

Cells were manually tracked using the MTrackJ plugin for ImageJ (http://rsb.info.nih.gov/ij). All cells on the bridge at the beginning of the experiment were tracked, excluding cells that died, to the end of the experiment or until the cells moved off the viewing bridge. Chemotactic index was calculated as described [5].

### siRNA knockdown

Cells were transfected with scrambled or gene-targeting siRNA using Lullaby transfection reagent (Oz Biosciences). Cells were transfected twice with 25nM siRNA, once every 48 hours, then assayed 48 hours after the second transfection. To assess gene knockdown, RNA was extracted from cells using an RNeasy mini kit (Qiagen) and cDNA synthesised from 0.5μg samples using a SuperScript III reverse transcriptase kit (Life Technologies) as specified by the manufacturer. qPCR was performed using PerfeCTa SYBR green FastMix (Quanta), with cDNA analysed in triplicate using a 7500 Fast Real time PCR system (Applied Biosciences), data was extracted using Applied Biosystems 7500 software version 2.0. QuantiTect primer kits (Qiagen) were used to assess gene expression of GAPDH, PPAP2A/LPP1, PPAP2B/LPP3 and PPAP2C/LPP2, with all gene expression normalised against GAPDH as an internal control using the comparative CT method [34].

### Mass Spectrometry

Cells were plated in a 6-well dish at a density of 1x10^5^ per well, then treated with siRNA. 24 hours after the final transfection of siRNA, old medium was removed and 2ml of fresh LPA or serum-containing RPMI added to each well. Media used for LPP knockdown experiments after 24 hours was serum-free RPMI with 10uM 18:0, 18:1 and 20:4 LPA added. Media used for LPP3 knockdown timecourse experiments was RPMI with 10% serum. 1ml of supernatant was collected at the indicated timepoints and a standard amont of synthetic 17:0 LPA added to the sample prior to butanol extraction [5]. LPA was analysed using a QExactive orbitrap coupled to a Dionex UltiMate 3000 LC system (Thermo Scientific). The LC parameters were as follows: 4μL of sample was injected onto a 1.7μm particle 100 x 2.1mm ID Waters Acquity CSH C18 column (Waters) which was kept at 50°C throughout the analysis. A gradient system of (A) aqueous 0.05% ammonium hydroxide and (B) 0.05% ammonium hydroxide/methanol was used, with a linear gradient of 0.3mL/min from 50% to 90% B over 7’ rising to 100% B within 1’ and maintaned for 2’. Thereafter, the column was returned to initial conditions and equilibrated for another 4 min. Lysophospholipids were analysed in electrospray negative ionization mode, with spray voltage 3 kV, capillary temperature 300°C, sheath gas flow rate 50, auxiliary gas flow rate 10. The resolution was set to 70,000, automatic gain control was set to 1 x 10^6^ with a maximum injection time of 200ms and the scan range was 300-700m/z. Peak areas were determined using MAVEN [35, 36] and concentrations normalised to the LPA(17:0) standard.

### Circular Invasion Assay

The circular invasion assay (CIA) was carried out as previously described [21, 33]. Briefly, an ibidi culture insert (ibidi) was adhered to the centre of a 35mm low μ-dish (ibidi), then 1x10^6^ WM1158 cells plated in the dish around the insert. The following day the insert and supernatant medium was removed and 200μl of Matrigel (Corning) diluted 1:1 with chilled PBS was added and left to set for 1 hour at 37°C, then 1ml serum-containing medium was added to the dish prior to imaging. If treated with DMSO, Ki16425 or HA130, cells were incubated for 15 minutes in medium containing the inhibitor/vehicle before removing the culture insert. DMSO/Ki16425/HA130 was also added to the Matrigel and medium added prior to imaging. The level of invasion after 24 hours was determined using ImageJ and calculated relative to the scrambled/DMSO treated controls.

### Spheroid Invasion Assay

WM852 cells were treated with siRNA as described above, then harvested 24 hours after the second hit of siRNA. The spheroid invasion assay was performed as per instructions using a Cultrex 96 well 3D Spheroid BME Cell Invasion Assay kit (Trevigen). Briefly, cells were resuspended in the Spheroid Formation Extracellular Matrix and 1x10^3^ cells were plated per well (in triplicate minimum per condition). The cells were left to form spheroids for three days, then Invasion Matrix was added to the invasion condition wells and culture medium was added to the control wells. Spheroids were imaged each day for six days following addition of the invasion matrix, using a brightfield microscope. Relative invasion was analysed using ImageJ by thresholding the spheroid area and measuring the number of pixels within that area. Each measurement per day was divided by the measurement taken on Day 1 (following addition of the invasion matrix) and calculated as a percentage of spheroid area on Day 1.

### Chemotaxis simulation model

The agent based model, modified from [19], simulated cells locally degrading a freely diffusing chemoattractant. Diffusion was simulated explicitly on a grid using a central differences approximation to the diffusion equation. The wells were modelled as large, finite and well-mixed reservoirs in contact with the grid. Cell movement was biased by the receptor occupancy difference across the cell, with receptor occupancy at a point **x** given by θ = c(**x**)/(c(**x**)+k_d_).

### Statistics

All graphs (excluding spider and rose plots) and statistics (excluding Rayleigh tests) were generated using GraphPad Prism 6. Spider plots were generated using Microsoft Excel. Rose plots were generated using the ‘circular’ package in R. Circular confidence intervals were denoted in red, which were calculated using the approach outlined by Upton & Fingleton [37]. The p-values for the circular range test were calculated using the formula stated in Mardia & Jupp [38], and the p-values for the Rayleigh Test were calculated using the method outlined in Mardia & Jupp [39].

## Conflicts of Interest

The authors declare no conflicts of interest.

## Acknowledgements

We are grateful to Dorothy Bennett, Elena Sviderskaya and the Wellcome Trust Functional Genomics Cell Bank for melanoma cells lines, Heather Spence, Ben Tyrrell and Niels van der Broek for technical help, Laura Machesky and Clelia Amato for helpful comments on the manuscript, BAIR for microscopy support and CRUK for funding.

## Supplementary Movie Legends

*Supplementary movie 1*

Representative timelapse movie of WM239A cells in a uniform serum environment. Cells in the “right cluster” display tracks in red and generally move to the right of the bridge over time, while those in the “left cluster” display tracks in yellow and generally move to the left.

*Supplementary movie 2*

Representative timelapse movie of WM239A cells in a uniform serum-free environment.

*Supplementary movie 3*

Representative timelapse movie of WM239A cells in a uniform serum environment in the presence of 10μM Ki16425.

*Supplementary movie 4*

Representative timelapse movie of WM239A cells, treated with either a scrambled control siRNA or siRNA targeting LPP3, in a uniform serum environment.

*Supplementary movie 5*

Representative timelapse movie of WM239A cells, treated with either DMSO as a control or 500nM HA130, in a uniform serum environment.

*Supplementary movie 6*

Representative timelapse movie of WM1158 cells, treated with either a scrambled control siRNA or siRNA targeting LPP3, in a circular invasion assay.

*Supplementary movie 7*

Representative timelapse movie of WM1158 cells, treated with either a DMSO control or 10μM Ki16425, in a circular invasion assay.

*Supplementary movie 8*

Representative timelapse movie of WM1158 cells, treated with either a DMSO control or 500nM HA130, in a circular invasion assay.

